# AI-enabled simultaneous phenotyping of leaf vein and stomatal traits uncovers independent genetic control in maize

**DOI:** 10.64898/2026.08.03.742415

**Authors:** Martina Sechi, Riccardo Porcedda, Martina Pallaoro, John N. Ferguson, Matteo Dell’Acqua, Andrea Vandin, Leonardo Caproni

## Abstract

**Background:** Leaves maintain hydraulic homeostasis during photosynthesis through the coordinated action of stomata, which regulate gas exchange and transpiration, and veins, which supply water to the leaf lamina. While functional links between stomatal and vascular traits are known in dicots, their potential genetic coordination in C4 crops remains poorly understood. We investigated the genetic architecture of these traits in maize using a Multi-parent Advanced Generation Inter-Cross (MAGIC) population and a low-cost, high-throughput phenotyping platform integrating leaf clearing, digital microscopy, artificial intelligence, and image analysis

**Results:** We phenotyped 285 recombinant inbred lines and the MAGIC founder lines, generating 8,072 images from 2,026 leaf samples taken from seedlings grown in controlled conditions. A YOLOv8-based model automatically detected stomata, while a custom and efficient image-processing pipeline quantified vein traits and stomatal spatial distribution patterns along cell bundles. This enabled simultaneous characterization of stomatal density, size, and distribution together with vein density, thickness, and bundle-associated spatial patterning.

Substantial phenotypic variation was observed among genotypes, with strong correlations between abaxial and adaxial traits but no significant correlations between stomatal and vein traits. QTL mapping identified 37 genomic regions associated with stomatal and vein traits, including loci containing known developmental regulators such as *stomatal density and distribution1* and *stomagen1*, as well as novel loci controlling stomatal spatial patterns, divergence between leaf surfaces and veins traits.

**Conclusions:** These results support independent genetic control of stomata and veins and decoupled contribution to water-use efficiency, providing a novel genetic framework to independently optimize leaf hydraulic capacity and gas exchange in target environments.

## Background

Climate change is threatening the agricultural yields of the world’s most important cereal crops due to rising global temperatures and constraints on water availability [1]. For maize, it is predicted that even a rise in global temperatures of 1.5°C to 2.0°C will significantly compromise yield potential and global supply chains [2]. Drought stress influences leaf transpiration as soil drying triggers loss of hydraulic conductivity, forcing stomata to close [3]. Although stomatal closure protects the plant against severe dehydration, it simultaneously restricts CO₂ uptake, leading to reduced carboxylation, impaired photosynthetic performance, and ultimate yield loss [4]. To enhance resilience to drought in maize, effective breeding approaches may look to harness genetic factors underpinning traits related to enhanced water-use efficiency, i.e. the ratio of carbon gain to water loss (Leakey et al. 2018). At the leaf level, this balance between carbon gain and water loss is structurally mediated by two leaf anatomical structures: stomata and veins [5]. Stomata mediate gaseous exchange between the leaf and the atmosphere and are key to optimising the balance between CO_2_ uptake and water loss [6], while veins ensure water supply to the leaf for transpiration and photosynthesis [7]. Stomatal conductance to water vapour (*g*_s_) is a function of stomatal features, e.g., stomatal density (SD) and size (SS) [8], as well as the degree of stomatal opening (aperture) [9]. *g*_s_ is pivotal in defining crop water-use and water-use efficiency (*WUE*) [10]. However, the rate of transpiration from the leaf is constrained not only by stomata, but also by the hydraulic capacity of the leaf. Minor veins provide the pathway for water delivery to evaporating tissues, and species with higher vein densities (VLA, minor veins length per unit area, herein VD) generally possess greater hydraulic capacity [5]. Consequently, stomatal and vein traits are often coordinated, with higher SD typically being accompanied by higher VD to maintain a balance between transpirational water loss and water supply [11,12]. The ratio of these two traits (SD/VD) can therefore provide a useful descriptor of the balance between evaporative demand and hydraulic supply within leaves [13–15], which varies both in response to evolutionary history and environmental conditions such as irradiance and water availability [13–15].

Maize is an amphistomatous species, meaning it has stomata on both leaf surfaces. Stomatal development and this abaxial-adaxial polarity are genetically controlled [16,17]. The link between stomatal patterning and water-use has motivated recent studies to manipulate key genetic modules in these developmental pathways in maize [18] and other grass species [19]. Typically, these efforts develop lines that show enhanced WUE. This is a particularly attractive approach in maize since it utilises a biochemical carbon concentrating mechanism via the C_4_ pathway, meaning photosynthesis should be less compromised by reduced *g*_s_ compared to C_3_ species [18]. Beyond stomata, vein density also exhibits phenotypic plasticity in response to environmental stresses such as elevated temperatures driving increased vein density and higher photosynthetic rates in maize [20]. While the physiological coordination between stomata and veins has been analysed in C_4_ species, including the impact on *WUE* [21], no studies, to date, investigated this putative coordination and their relative leaf polarity in maize.

Different phenotyping approaches have been developed to investigate stomatal patterning alone. These start from low-cost but time-consuming methods, like the combination of nail varnish and tape on the leaf to produce micrographs through light microscopy, often obtaining high-definition images [22,23], and extend through to high-throughput methods that utilise digital handheld microscopes directly on the leaf [24] or optical topometry [25,26] where high numbers of epidermal micrographs can be produced rapidly without sample preparation. Often, these methods are supported by AI-based models allowing the processing of vast numbers of images with optimum stomatal detection. However, these methods cannot detect veins, since they become visible only after clearing the tissue [20,21]. Whilst models have been developed to analyse the spatial distribution patterns of stomata across different species, including maize [27–29], a key phenotyping bottleneck is the availability of methods that can accurately quantify both stomatal and vein traits on the same leaf sample.

Leaf anatomical traits are complex traits, regulated by different genomic regions. Genetic mapping has been widely employed to dissect complex traits into quantitative trait loci (QTL), and to identify allelic variations that can provide opportunities for breeding to develop new climate-resilient varieties [30–33]. In crops, bridging the phenotype-genotype gap remains challenging. To overcome it, genetic material that provides high statistical power, while minimising the high number of genotypes used, should be employed [34]. Diversity panels present high genetic diversity and historical accumulation of recombination events, however mapping populations, crossing two or more parental lines, offer higher precision in detecting QTL due to the known pedigree and low genetic structure [34]. In maize, the Multiple parent Advanced Generation Inter-Cross (MAGIC) population, developed in 2015 [35], has become a promising tool to detect genetic determinants of traits of agronomic interests such as photosynthetic efficiency [36,37], enabling high genetic diversity and high recombination density.

In this study, we aimed to dissect the genetic determinants of key structural and hydraulic leaf traits that influence WUE by combining the statistical power of the MAGIC maize population with a novel, low-cost, high-throughput phenotyping approach for characterising both stomatal and vein traits on the same leaf sample. Our AI-enabled phenotyping approach allowed us to characterise established and novel stomatal and vein traits across more unique genotypes than has ever previously been demonstrated. Downstream QTL mapping demonstrated the presence of pleiotropic genetic regions influencing both stomatal and vein traits, but also QTL influencing these complex structures in isolation. From a crop improvement perspective, the discovery of QTL that influence vein traits independently of stomatal patterning is particularly important as it suggests that breeding may be able to modify hydraulic capacity without simultaneously altering stomatal characteristics, thereby permitting independent optimisation of water transport and gas exchange according to the target environment.

## Materials and methods

### Genetic materials

In this study we used 285 Recombinant Inbred Lines (RILs) and the founders of the MAGIC maize population [35] (Supplementary Table S1). In brief, the population was developed starting from 8 inbred lines (A632, B73, B96, F7, H99, HP301, Mo17, and W153R), following a half-diallelic breeding design. A ninth line (CML91) was introduced into the population as a 2-way hybrid (B73xCML91) to account for failures in crossing of B96xHP301.

### Greenhouse experiment

The study was conducted on tissues from seedlings grown in a greenhouse trial, carried out from October to December 2024 at the University of Milan (Milan, Italy). We used an augmented block design developed with R/agricolae [38]. It consists of four batches and four blocks per batch. Six out of nine parental lines, namely A632, B73, CML91, H99, HP301, and W153R, were used as controls and replicated four times in each batch (*i.e.* one replicate per block). In each batch, we sowed 18 seeds per line, for 72 RILs (included 3 founders) and the 6 control genotypes (i.e. six out of the eight MAGIC founders). Seed were placed at 2 cm of depth, in six-wells trays (164 x 150 x 58 mm, F.P. PLAST SRL, Scandiano (RE), Italy) containing S.Q.10 substrate (peat, sand, compost; Vigorplant Italia S.r.l., Fombio, Italy). Plants were grown under a long-day photoperiod (16 h light/8 h dark) with temperature set to 25°C and with photon fluence of 270 mmol m^-2^ s^-1^. Seven days after sowing (DAS), plants were fertilized with a commercial liquid fertilizer (COMPO Concime per Piante Verdi®, COMPO Italia S.r.l., Italy; NPK 7.5–3–6, supplemented with Fe) applied twice a week during second week after sowing. We monitored seed germination and removed excess seedlings to obtain 6 replicates per genotype within a single tray.

### High-throughput phenotyping of maize leaves

Fourteen DAS, we cut the second fully expanded leaf from six replicates per genotype. An 8-cm leaf segment was excised starting from the leaf apex. The cuts were submerged in a 15 ml falcon tube containing a solution of 80% [v/v] ethanol and incubated at room temperature for a week and then stored at 8°C for the following three months. Five days before imaging acquisition, the solution was replaced with fresh ethanol 80% [v/v] solution to allow the leaves to promote additional bleaching, and to improve the visibility of veins and stomata. For each replicate, the leaf segment was mounted on a glass slide; the distal and proximal 2 cm portions were removed to retain only the central region for imaging. This section was placed between two microscope slides (VWR, Avantor) and then over a light source to facilitate the visualization. Images were acquired using a handheld digital microscope (DinoLite AM4115T – JV, AnMo Electronics Corporation, New Taipei City, Taiwan). For each sample, two images were acquired from both the abaxial and of the adaxial side using the midrib as a reference point (Fig. 1A). The images were collected with a 220x magnification, corresponding to a field of view of 2.39 mm^2^. Images were processed using a DinoCapture 2.0 software with a resolution of 1280 x 1024 pixels.

**Figure 1.**
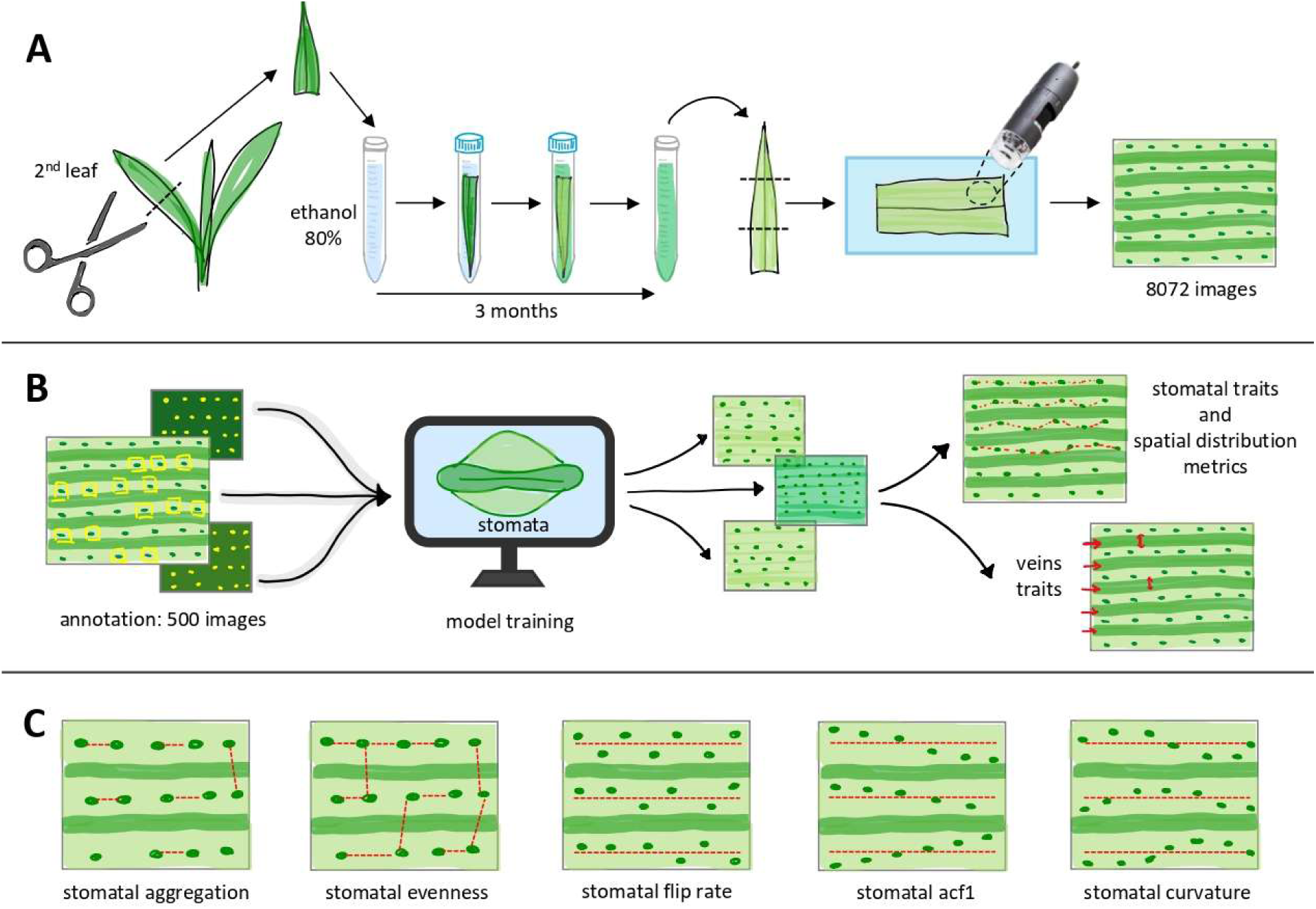
A) Phenotyping with ethanol 80% solution and handheld digital microscope. B) Image annotation, YOLOv8 training and detection of stomata on images. C) Graphical explanation of spatial distribution patterns of stomata.

### Dataset

In order to proceed with the training of a stomatal detection model, we first randomly selected 500 images and uploaded to Roboflow (Roboflow Inc. 2025) to detect and annotate the stomata in each image, employing the bounding box annotation tool (Supplementary Fig. S1). After dividing the images into training (350), validation (101) and test (49) set, we generated two augmented versions of each training image in Roboflow, increasing the training set to 1,050 images. We applied further augmentation dynamically during training, including four-image mosaic composition, horizontal flipping with a probability of 0.5, translations of up to 10%, scaling between 50% and 150%, and variation in hue, saturation, and brightness. We also applied low-probability blur, median blur, grayscale conversion, and contrast-limited adaptive histogram equalization (CLAHE; probability 0.01 each). The entire process is illustrated in the Fig. 1B. Finally, we exported the annotated images via API to train an Oriented Bounding Box (OBB) detector using the Ultralytics YOLOv8 architecture [39] in Python.

### Fine-tuning a pretrained model

Training a model starting from random initialization of the weights requires time and a non-trivial effort of annotation of a large number of images. For this reason, we decided to apply *transfer* learning, i.e. using a pre-trained model already capable of detecting objects from other domains and adapt it with few epochs of fine tuning on a small set of annotated images of the domain of interest [40]. We adopted the YOLOv8n-obb weights (transferring 391 out of 397 compatible weight tensors) and the detection head was adapted from the original 15 classes to the single *stomata* class. The model was subsequently fine-tuned on the annotated images. The choice of OBB version of YOLOv8 is motivated by the need for the extraction of the bounding box area in rotated images (i.e. the right estimate for the detected stomatal area).

Model performance was evaluated on the test set using standard object-detection metrics, including Precision, Recall, mAP@0.5, mAP@0.5:0.95 and F1 score. For YOLOv8-OBB, these metrics were calculated using probabilistic Intersection over union (IoU, i.e. ProbIoU). ProbIoU represents each oriented box as a two-dimensional Gaussian distribution and evaluates similarity in position, dimensions, and orientation through a Hellinger-distance-based measure. This provides a smooth comparison that is particularly suitable for rotated or approximately elliptical objects, for which small differences in box angle may not be meaningful.

For each validated detection, the model returned the centre coordinates (c_x_, c_y_), width (w), height (h), orientation angle (θ), and confidence score (s). Stomatal area was estimated as w*h.

### Remarks on model calibration

As it is usually done in the field of machine learning, we first selected the confidence threshold by choosing the one that maximized the F1 score on the validation set. As it will be shown in the Results, this led to an overestimation of the count of stomata in each leaf image. For this reason, and since the downstream analyses also depend on accurate stomatal counts, we defined a second, count-balanced operating threshold using the validation set. For each candidate confidence threshold, we calculate for each image the difference between the count of detected stomata and the count of annotated stomata.

In symbols, Δ = *N*_*predicted*_ − *N*_*annotated*_ and the threshold minimizing |*mean*(Δ)| + *SD*(Δ) was selected.

### Vein traits detection

We developed an efficient Python pipeline of image processing to extract vein traits from each image. Images were converted to grayscale, and the dominant vein angle was estimated using an image structure tensor. Images and their corresponding stomatal coordinates were then rotated by the opposite angle to align the main veins horizontally, followed by contrast normalization (stretching between 2^nd^ and 98^th^ percentiles), adaptive gamma correction, and binarization via Otsu thresholding. Main vein bands were identified from the row-wise dark-pixel profile of the aligned binary mask (the same procedure was applied with white-pixel profile for bundle bands). From these detected structures, the pipeline extracted the longitudinal vein thickness (LVT), bundles thickness (BT), longitudinal vein density (LVD), that was estimated as the total length of all detected veins divided by the total analysed image area, expressed in px^-1^.

### Calculation of stomatal spatial distribution patterns

The simultaneous analysis of veins and stomata, which is the core of this work, allowed us to study the spatial distribution of stomata inside bundles. We calculated the spatial distribution patterns (SDP) of stomata using two complementary approaches (Fig. 1C and Supplementary file S1). Firstly, we computed global descriptors, recently developed and applied on different tree species [27]. These included the stomatal aggregation index (R), defined as the ratio between the observed mean nearest-neighbour distance and the expected distance under a random Poisson pattern, distinguishing clustered from regular patterns. Then, stomatal evenness (E) was derived from a minimum spanning tree connecting all stomatal centres to evaluate the homogeneity of distances between neighbouring stomata.

Secondly, we computed local descriptors within cell bundles by relating stomatal positions directly to the local venation architecture. Images were converted to grayscale, and the dominant venation angle was estimated using an image structure tensor. Stomata were assigned to specific bundles based on their vertical coordinates within these defined bands. Within each bundle, we fitted a constrained linear regression line with a fixed slope matching the dominant venation angle to establish the central axis. We then calculated the signed distance (residual height) of each stoma relative to this baseline, ordering stomata along the bundle’s longitudinal axis to generate a bundle-specific height profile.

From the resulting ordered height profiles along each bundle, three local patterns were calculated and averaged per leaf image. Flip rate shows the frequency of direction changes around the bundle axis, first-lag autocorrelation (ACF1) quantifies whether a stoma’s height depends on the preceding stoma along the bundle and curvature is computed from second discrete differences of the height profile to quantify local spatial irregularity. In Supplementary Fig. S2, we show how different combinations of these descriptors match typical observed spatial distribution patterns of stomata inside bundles. In Supplementary file S1 we give further details on the definition of the spatial distribution metrics mentioned above.

### Trait processing and statistical analyses

After the training, we processed the resulting data exclusively within the R programming environment, if not stated otherwise. For abaxial and adaxial side we calculated the stomatal density (SD) as the number of stomata/unit area (mm^-2^). We calculated the area of the OBB, as a proxy of stomatal size (SS), as A=b*h and converted to µm^2^. We checked the correlation between the two leaf sides through Person correlation and tested the differences between the two leaf sides with the Wilcoxon rank sum test. Also, we checked the distribution and normality of the data. We employed the Kruskal-Wallis rank sum test to evaluate the experimental design effect on the entire dataset and then on the founder lines. Using the *identify_outliers()* function of rstatix, we detected and filtered out the extreme outliers for RILs and founders within each genotype and leaf side. We used this statistical approach for the SDP too.

For all the stomatal traits and SDP, we derived the Best Linear Unbiased Predictors (BLUPs) fitting a Linear Mixed-Effect Model (LMM) through the *lme()* function of lme4 [41]. The model included leaf side as fixed effect and genotype as random effect. We added a genotype-by-leaf side interaction as random effect and a hierarchical random effect structure of plant replicate within block, which was nested within batch. We calculated the BLUPs for each side by integrating four model components (equation 1): the population intercept (abaxial as intercept), the main genetic effect for each genotype, the specific interaction effect for that genotype on that specific leaf side, the fixed effect of leaf side (abaxial side as the reference). We extracted the components using *ranef()* and *fixef()* function of the lme4.

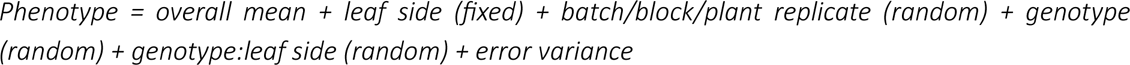

For each image, we derived the number and the median thickness of veins and cell bundles. Then, we converted the thickness in µm and the density in 1/mm. For all the vein traits, we used the same statistical approach used for the stomatal traits. We detected the extreme outliers for vein number, removing all the information for those images.

We derived the BLUPs for all the traits fitting a LMM used for the stomatal traits, without including the interaction between genotype and leaf side. Then we calculated the BLUPs for each genotype combining the estimated fixed effect with the predicted random effect (equation 2).

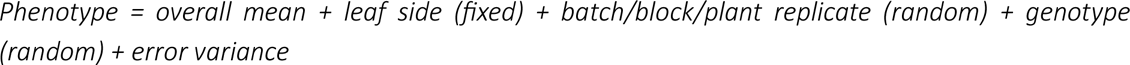

From the extracted BLUPs for each genotype we calculated the total SD for each leaf as the sum of SD_abaxial_ + SD_adaxial_, the ratio SD_adaxial_/SD_abaxial_, the percentage of the leaf area covered by stomata (SA) as SD*SS and the ratio SD_abaxial_/ VD_abaxial_. We calculated the ratio adaxial/abaxial for all the stomatal traits (additional details in Supplementary Table S2). For each stomatal and vascular traits, we calculated the broad-sense heritability (H^2^_B_) using the function *H2()* of heritable. Then, we checked the distribution of the BLUPs using boxplot combined with jitter plot. We assessed the phenotypic relationship by calculating a Pearson correlation matrix on all the traits using the function *corrplot.mixed()* of the corrplot. We performed a Principal Component Analysis (PCA) with stats and ggfortify to explore the structure of phenotypic diversity within the MAGIC population, using the eight founder lines as distinct reference genotypes.

### QTL mapping and identification of candidate genes

We employed 79,557 SNPs, mapped on V4 genome reference version [42], and obtained through the Single Primer Enrichment Technology (SPET) with a custom set of probes [43]. For qtl mapping we used functions implemented in r/qtl2 [44]. Genotype probabilities were calculated using a Hidden Markov Model (HMM) with the function *calc_genoprob()*. The kinship was derived through the “Leave One Chromosome Out” (LOCO) method. Logarithm of odds (LOD) scores were estimated using a Linear Mixed Model that incorporated the kinship with the function *scan1()*.

Peaks were identified using a Bonferroni correction for multiple testing and calculated the significance threshold with alpha as 0.05. QTL peaks were identified with the function *find_peaks()*, specifying drop and peak drop of 1.5. We used a maximum QTL span threshold of 10 Mbp to identify candidate genes, in genomic regions with relatively strong statistical support, only [36,37]. At these selected QTL, founder’s haplotype coefficients at were estimated by fitting a generalized linear model using the function *scan1blup()*. The gene models in the selected QTL spans were derived from the B73 RefGen_V4 annotation and used to search the *Arabidopsis thaliana* best-hit orthologs using The Arabidopsis Information Resource (TAIR 11, [45]). Where available, the functional descriptions and Gene Ontology (GO) terms for these orthologs were retrieved.

## Results

### Accelerating phenotyping through high-throughput imaging pipeline

We grew 285 RILs and 9 founders of the MAGIC maize population under controlled greenhouse conditions to capture variation of stomatal and vein traits. Fourteen DAS, we successfully sampled 274 RILs (96% of the population) and all founders, yielding a total of 2026 leaf samples (90,3% of the theoretical total number with at least 2 replicates per genotype). Leaf cuts were treated in 80% (v/v) ethanol prior to imaging. Leaf images were acquired on both abaxial and adaxial surfaces, resulting in a total of 8072 images. We fine-tuned a pretrained YOLOv8n-obb model with a maximum of 200 epochs and early stopping after 30 epochs without improvement. The model achieved rapid convergence, stopping at epoch 100 with the optimal weights retained from epoch 70.

The confidence threshold was selected as the value that maximized the F1 score on the validation set; this threshold was 0.3704, yielding an F1 score of 90.3%. With this threshold and the ProbIoU-based evaluation, we obtain a mAP@50 of 96.8% and a mAP@0.50:0.95 of 77.2%, In addition, we evaluated localization using exact polygon IoU, defined as the geometric intersection area divided by the union area of the predicted and annotated oriented polygons. Predictions and annotations were assigned one-to-one using Hungarian matching and were considered true positives at polygon IoU ≥ 0.50. This led to an identification of 4,043 true positives, 643 false positives and 322 false negatives, which translates to 86.3% of precision, 92.6% of recall and 89.3% of F1 score.

The mean polygon IoU among true-positive matches is 79.1%. When missed annotations are assigned an IoU of zero, the mean IoU over all ground-truth stomata is 73.2%.

We proceeded with the count-based calibration to identify a confidence threshold of 0.475. At this threshold, the model detected 7,552 stomata across the validation set, compared with 7,554 annotations, corresponding to an aggregate count bias of −0.03%, a median per-image difference of zero, and a mean absolute error of 7.33 stomata per image. At the F1-selected confidence threshold of 0.3704, the model detected 7,863 stomata in the validation set, that is to say 309 additional detections with respect to the annotated stomata. Exact-polygon detection test performances remained similar: with respect to the results with F1-selected confidence threshold,

Across the entire image dataset, our automated pipeline identified only three images without stomata, which we subsequently excluded from the downstream analysis. Furthermore, we removed a single RIL due to sub-optimal image quality, which prevented reliable automated feature extraction. Intra genotypic variance was exceptionally low for all the traits, with extreme outliers accounting for only 0.5%-1% of the observations in the entire dataset.

### Population-Level Variation in Stomatal and Vein Traits

The raw trait values highlighted high and positive correlation between abaxial and adaxial traits (Supplementary Fig. S5). These correlations were higher when BLUPs were used (Supplementary Fig. S6): for SD (r=0.82, p<0.001), and SS (r=0.97, p<0.001) was relatively high, while for SDP lower (from r=0.65 to 0.85). For all vein traits the correlation was consistently high (r=1, p<0.001).

For all the stomatal and vein traits broad-sense heritability (H_B_^2^) ranged between 0.62 to 0.95 (Cullis, Supplementary Table S3). The distribution of the BLUPs of all the traits (Supplementary Fig. S7) shows a wide phenotypic variation at population level, reflecting the phenotypic variation at founders’ level (Fig. 2A-F).

**Figure 2.**
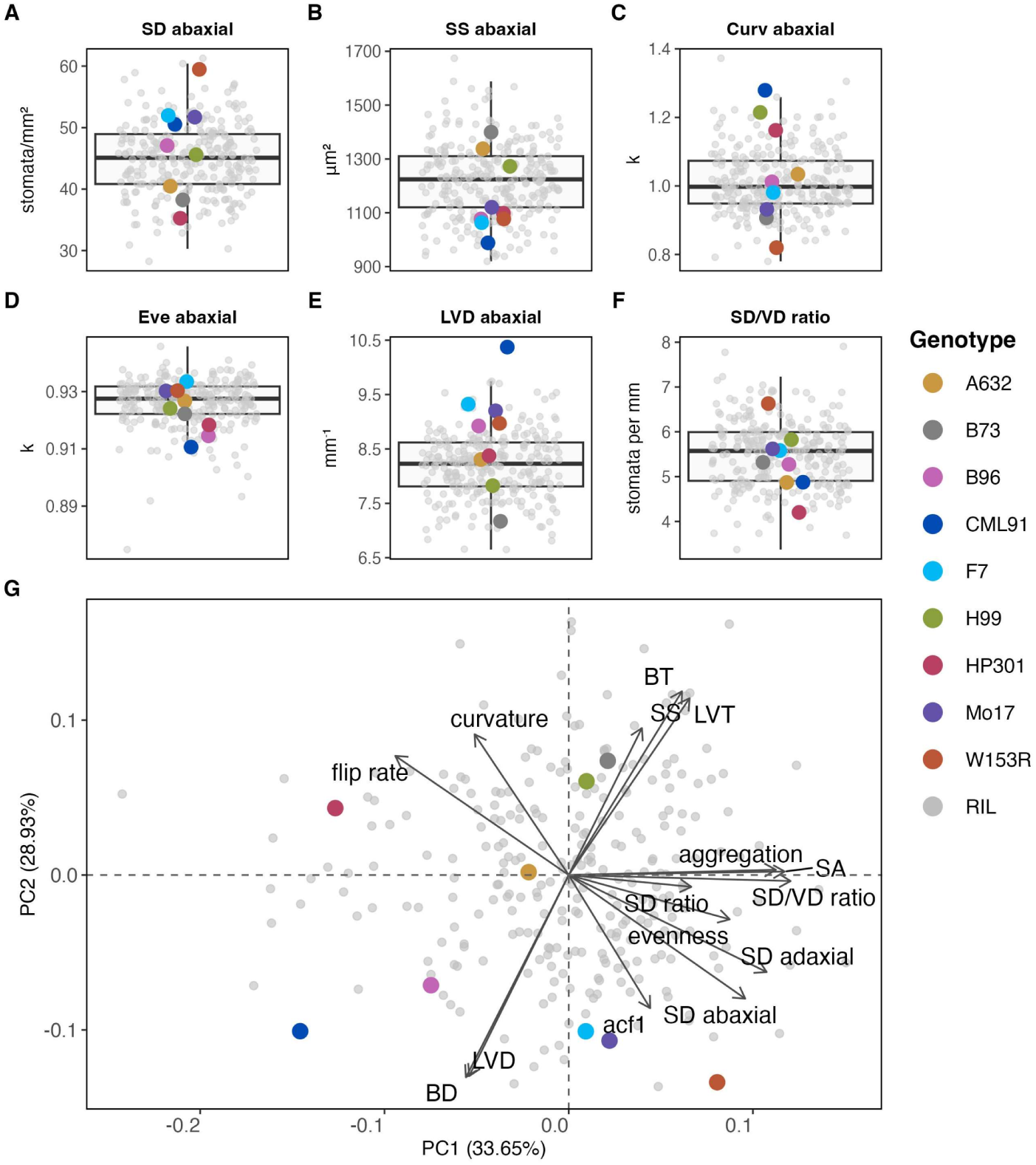
A-F) Boxplots of BLUPs on six traits of 273 RILs and 9 founder lines. RILs are grey dots while founder lines are coloured. G) PCA of BLUPs of 273 RIL and 9 founder lines.

The boxplots show continuous distributions for all the traits, but also the presence of some RIL positively or negatively exceeding the phenotypic expression of the founders. SD (Fig. 2A) shows an overall variability of 3-fold, also observed among the founders, while the SS (Fig. 2B) shows a of 4-fold and 3-fold among the RILS and founders, respectively. Curvature and aggregation (Fig. 2C and D) have an overall variability of 3-fold and 2-fold, respectively, that reflects the variability among the founders. LVD (Fig. 2E) shows a high overall variability of 4-fold, which is observed also among the founders. The ratio SD/VD on the abaxial leaf side (Fig. 2F) presents an overall variability of 4-fold, while a 3-fold variability among the founders.

To evaluate the phenotypic diversity of the population on all leaf anatomical traits we performed principal component analysis (PCA) (Fig. 2G), with founder lines as reference. In the PCA, the first two PCs explain 62.58 % of the total phenotypic variance. Specifically, SD, SA, SD/VD ratio, aggregation, evenness and ACF1 contribute positively to the variation along the PC1, whereas flip rate and curvature project in the opposite direction. On the other hand, vein and bundle thickness (LVT, BT) alongside SS show a positive contribution to the diversity along PC2, while vein and bundle density (LVD, BD) align in the negative direction.

Correlation analysis of BLUPs (Supplementary Fig. S6) revealed several significant relationships among the measured traits. SD was negatively correlated with SS (r = 0.31; Abaxial). Among the SDP traits, SD was negatively correlated with flip rate (r = -0.52) and curvature (r = -0.28) but showed no significant correlation with ACF1. In contrast, SD with positively correlated with aggregation (r = 0.49) and evenness (r = 0.39). The SD/VD ratio was positively correlated with SA (r = 0.66) and aggregation (r = 0.41) and negatively correlated with flip rate (r =-0.39). SD showed weak positive correlations with LVD (r = 0.21) and BD (r = 0.20), but no significant correlation with thickness. Conversely, SS was moderate negatively correlated with LVD (r =-0.47) and BD (r =-0.49) and positively correlated with LVT (r = 0.53) and BT (r =0.49). Vein and bundle-related traits were positively correlated with one another, whereas density and thickness traits were strongly negatively correlated, with LVD versus LVT (r =-0.82) and BD versus BT (r =-0.89). Finally, the SD/VD ratio was more strongly correlated with SD (r = 0.84) than with VD (r =-0.35), indicating that variation in the ratio was driven primarily by SD rather than VD (Supplementary Fig. S6).

### Genetic architecture of stomatal and vein traits

Across the genome, we identified 89 significant signals (Supplementary Table S4), 56 of which consistently defined 37 potential QTL spanning physical intervals less than 10 Mbp (Fig. 3A; Supplementary Table S5). These QTL mapped to 29 distinct genomic regions, with the identified signals covering an average confidence interval of 4.4 Mbp. Signals were found on all chromosomes, with most of the hits on chromosome 9. Regarding stomatal traits (SD, SS, SA and SDP), 53.5% of the identified QTL mapped to the abaxial leaf side, whereas 46.5% mapped to the adaxial side. SD shares just one region between the two leaf sides on chromosome 6, while SS shares 3 distinct QTL on 3 distinct chromosomes. QTL controlling SD map to distinct regions from those of SPD, except on chromosome 4 and 6. QTL on chromosome 4 is also shared with SS, which co-localise on chromosome 9 with SDP. SA shares two region on chromosome 4 and 6 with SD and SS. Vein traits do not share any QTL with SD, while they share 3 signals on chromosome 1,5 and 9 with SS. Also, vein traits define distinct QTL from those of SPD, except one on chromosome 9. Ratio SD/VD maps to one QTL on chromosome 6, co-localised with SD.

**Figure 3.**
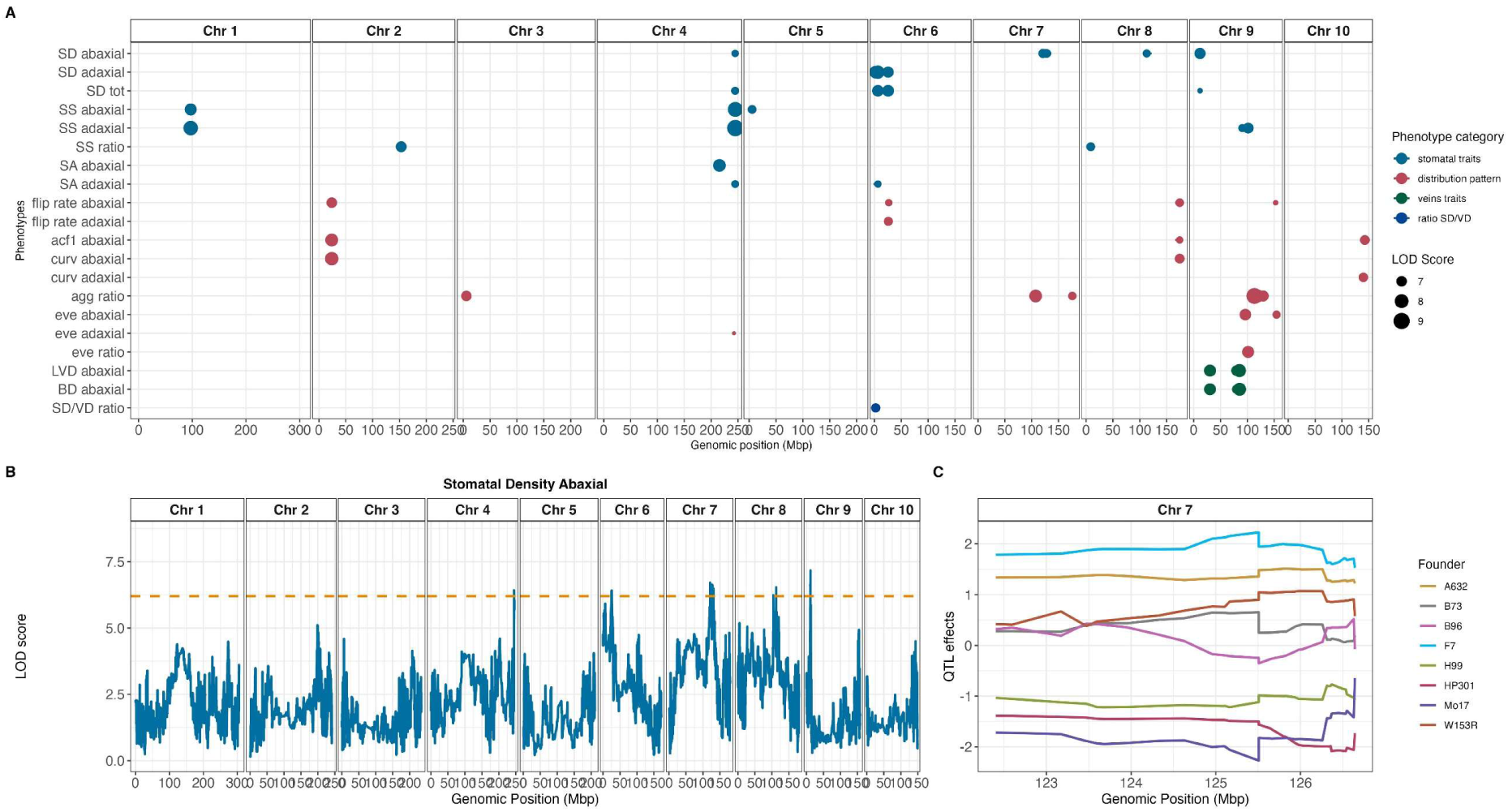
A) Summary plot of QTL peaks detected with QTL mapping for all the traits. B) LOD score plot of stomatal density abaxial with significant thresholds. C) Plot with founder effect on the QTL detected on chromosome 7.

These 37 QTL harbour 2163 unique gene models (Supplementary Table S6). Among these genes, we could identify 10 known to be involved in stomata development in plants (Table 1). Two out of ten were already characterised in maize: *stomatal density and distribution 1 (SDD1)* and *stomagen1*. The two QTL, harbouring these genes, were mapped with abaxial SD and SDP-derived traits (Fig. 3A) on chromosome 7 and 8, respectively. Some of the traits, like abaxial SD (Fig. 3B), map multiple QTL, including the chromosome 7 region at 126 Mbp where *SDD1* is present. At this region the founder haplotype effects of HP301, Mo17 and H99 drive a negative trait expression, meaning that RILs carrying these haplotypes at the locus, show lower abaxial SD (Fig. 3C). On the other hand, the F7 haplotype exerts a opposite effect, contributing to higher abaxial SD. A similar founder haplotype effect pattern could also be observed for the chromosome 8 QTL (174Mbp), harbouring *stomagen1*. The remaining 8 candidates (Table 1) were mapped with abaxial SD, SDP and LVD, and are the best-hit orthologs of *A. thaliana*’s *phosphatase 2C5 (P2C5)*, *mitogen-activated protein kinase 4 (MKK4)*, *cyclin D4*, *cell division control 6B (CDC6B)*, *epidermal patterning factor 2 (EPF2)*, *galacturonosyltransferase 10 (GAUT10)*, *galacturonosyltransferase 11 (GAUT11)* and *ERECTA-like 2 (ERL2)*.

**Table 1.**
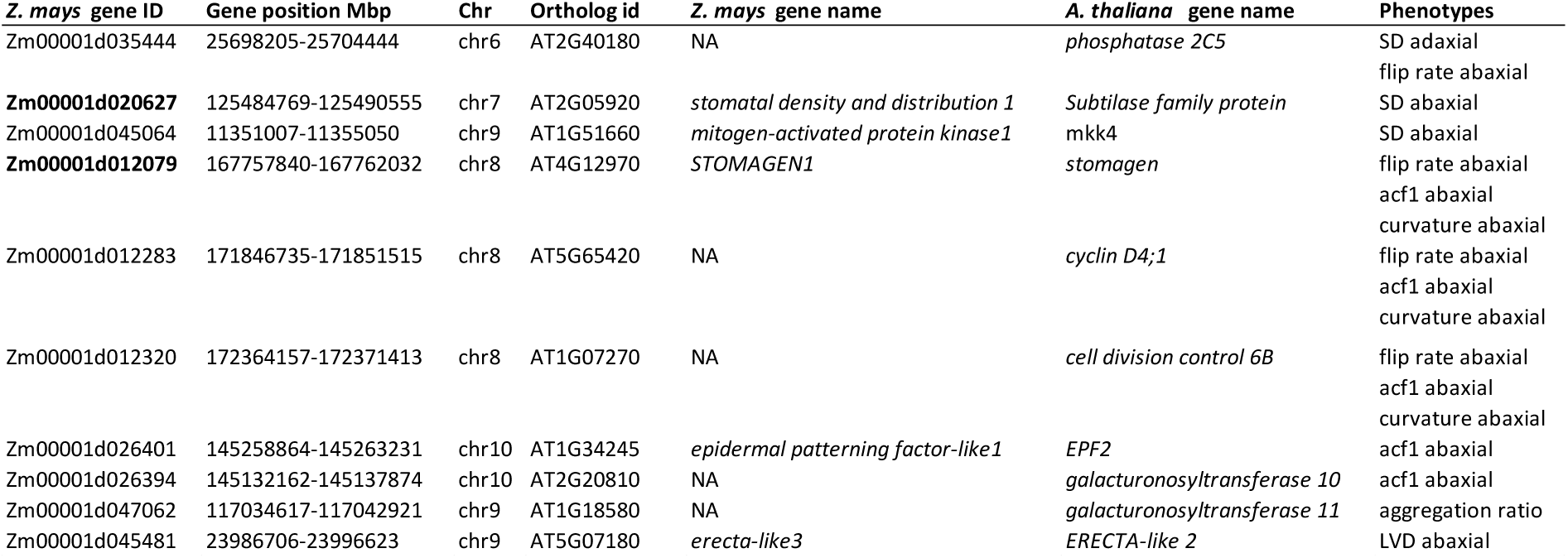
List of candidate genes with putative function in stomatal development. C.I. stands for Confidence interval. The genomic positions of Peak and CI are reported in Mbp. *Z. mays* gene IDs highlighted in bold are those known to be involved in stomatal development in maize.

## Discussion

Recent work contributed to the understanding of the genetics underpinning stomatal patterning in maize and other C4 grasses [25,26,29,46]. Concurrently, physiological coordination between stomata and veins related to WUE has recently been described in sorghum [21]. However, the underlying genetics of this coordination has not yet been explored specifically through quantitative genetics. In this study, we implemented a high-throughput pipeline combining a low-cost imaging approach with AI-based and computational analyses to derive both stomatal and vein traits at the same time. This allowed us to explore unstudied spatial distribution patterns of stomata and map the genetic determinants of the leaf transpiration and water supply. To achieve these objectives, we grew a MAGIC maize population in controlled conditions using a fully replicated experimental design. We successfully sampled and processed 2026 leaves and 8072 images for both abaxial and adaxial leaf sides with an investment of about 300 man-hours. To our knowledge, this is the first study to phenotype stomatal and vein traits on the same biological samples and perform QTL mapping for these traits on both leaf surfaces.

The application of YOLO model to automatically detect stomata on images has been already tested across several species [24,28]. By fine-tuning a pre-trained YOLO model on a small annotated dataset, we maintained model simplicity while significantly reducing training data requirements. Combining AI-based stomatal detection with Python-based vein extraction allowed us to process a large image dataset, offering a user-friendly approach to accelerate phenotyping of these traits on the same leaf. However, our pipeline could not capture finer stomatal features, such as aperture or guard cell metrics. Integrating our sample preparation with low-cost with higher-magnification devices could enable characterization of these fine-grained morphometrics in future work.

Taking and processing images on both abaxial and adaxial leaf side cleared by ethanol solution allowed us to type a diversity of stomatal traits, their distribution patterns as well as vein traits. Results showed that, at population level, SD is higher on abaxial than adaxial side, which is in line with the amphistomous nature of maize [47] and reflects the higher evaporative demand experienced by the adaxial leaf surface [47]. The leaf surface polarity decreases considering the stomatal size, vein and cell bundles density and thickness. The patterns related to the linearity of the stomata along the cell bundles (flip rate, ACF1 and curvature) and evenness and aggregation show a moderate difference between the two leaf sides, showing a higher correlation to SD rather than the SS. All stomatal and vein traits showed moderate to high broad-sense heritability (H_B_^2^= 0.62 to 0.95, Supplementary Table S3), suggesting that the observed phenotypic diversity across the traits is mainly driven by the genetic factors. However, these estimates should be interpreted with caution, as they were obtained from a single experiment conducted under controlled environmental conditions. Given that stomatal development is known to be influenced by environmental factors [48,49], the relative contribution of genetic and environmental effects may differ under field conditions.

For all the traits, we derived BLUPs enabling the investigation of phenotypic correlation among traits. SD was negatively correlated to SS, confirming the trend highlighted by different studies in maize [25,50,51]. In contrast, SS is negatively correlated with VD and positively with VT, reflecting the anatomical and spatial organisation of stomata. The anatomical coordination between stomata and veins has already been characterized across C3 species [12] and within C4 species [21]. At the physiological level, this coordination allows the plant to establish a trade-off between water supply and evaporative demand. Indeed, different species that display high SD and high VD also show relatively high stomatal conductance (g_s_) [5]. In C3 and C4 species, g_s_ is correlated to water use efficiency (WUE) [10]. Studies on sorghum and maize have shown that reducing g_s_ through lower SD can improve WUE [18,19], while a recent study on maize has revealed a correlation between high vein density and the improved photosynthetic performance [20].

Building on these studies, we hypothesize that selection, either natural, human or a combination of both, has shaped the coordination between stomata and veins to balance water demand and supply across different environments. In our mapping population, we did not observe such a correlation. The genetic constitution of the population, with its relatively high level of recombination, enabled the independent segregation of loci controlling vein and stomatal traits.

The ratio between SD and VD reflects the coordination between transpiration through stomata and water supplied by veins in the leaf [11,12]. Our results displayed that the SD/VD ratio high variation is highly dependent on SD rather than VD, suggesting that stomatal density is the primary driver of the variation in this anatomical ratio and may therefore play a more predominant role in modulating maize WUE. Consequently, we hypothesise that reducing the SD, achieving a lower SD/VD ratio and maintaining a higher vein supply relative to stomatal count, could represent an efficient breeding strategy to develop new maize varieties resilient to drier environments.

Considering the genomic contribution of the founders, in terms of haplotypes to the diversity at population level (Fig. 2A-B-E-F-G), we observe that F7, Mo17 and W153R contribute to higher SD, indicating a potential decreased WUE and, consequently, photosynthesis [37]. All these three lines can be classified as from temperate origin, as most of the founders, except B96 and CML91. Indeed, these lines display similar combinations of stomatal and vein trait expressions. Among the temperate lines HP301 also has a low SD/VD ratio, appearing to be the line with the most conservative strategy in maintaining the balance between the water supply and the evaporative demand.

In this research, we successfully applied aggregation and evenness previously used for tree species, to successfully type spatial distribution of stomata in maize [27]. In addition, we evaluated flip rate, ACF1, and curvature to further quantify stomatal spatial patterns along cell bundles. The results suggest that when SD increases, stomata tend to be distributed more evenly on the leaf surface. However, their arrangement does not follow a linear pattern along the cell bundles; instead, stomata form irregular zig-zag or curved spatial configuration. On the contrary, we did not find any correlation between the vein traits and the spatial distribution patterns, hypothesising that the correlation between SD and the patterns are purely regulated by anatomical constraints instead of physiological reasons. To confirm this hypothesis, photo-physiological phenotyping to contrast distinct haplotypes based on these distribution traits will be required.

We detected 37 QTL having relatively high statistical support (Fig. 3A). Consistent with the lack of correlation observed between SD and vein traits, these traits map to distinct regions. However, few veins-related QTL co-localise with loci associated with SS and SDP, suggesting potential pleiotropic effects or tight genetic linkage underlying the regulation of stomatal morphology, patterning and vascular architecture. We identified 2163 unique gene models (Supplementary Table S6). Among these genes we reported those already or putatively associated with the phenotypes. For SD, we identified different leaf side-specific QTL, suggesting that adaxial and abaxial stomatal development may be controlled independently [52]. The QTL identified for SD appear to be different from those detected with one among the maize-NAM subpopulation (B73XMS71) [25], except the chromosome 7 QTL. Despite sharing the B73 founder, the NAM subpopulation was evaluated in the field on adult plants; these distinct conditions likely explain the differences detected for this trait. On the same QTL, we found that F7 harbours the positive allele effect on abaxial SD, suggesting that this haplotype may be involved in the increase of SD on the abaxial leaf side, potentially resulting in a reduced WUE [18,19]. This QTL harbours *SDD1*; previous evidence shows that its overexpression reduces stomatal density and conductance, without compromising photosynthesis [25]. For SD we also found the arabidopsis’ best-hit orthologs of *P2C5* (chr 6, 26 Mbp) and *MKK4* (chr 9, 12 Mbp), known to be involved in stomatal aperture [53,54]. However, these genes are not characterised yet in maize, therefore we suggest them as novel putative candidate genes for abaxial SD.

SD can be seen as complex trait, and therefore, its decomposition into simpler and easier to measure components, can help identifying their genetic bases. Indeed, the few co-localised QTL among SD, SS, SA and SDP suggests a partial independent genetic control. We could observe the same condition for flip rate, ACF1 and curvature vs. aggregation and evenness, where only a single QTL is shared among the traits. For abaxial flip rate we found again the arabidopsis’s orthologs *P2C5*, while for abaxial flip rate, ACF1 and curvature we identified *stomagen1*, that in maize mutants is associated to lower SD and index, but also associated to reduced net photosynthetic rate, transpiration rate, g_s_ but increased WUE [55]. On this QTL on chromosome 8, HP301, H99 and Mo17 harbour a positive allele effect, resulting in an increased irregularity of the stomata along the cell bundles, that we found to be negative correlated to SD.

For abaxial flip rate, ACF1, and curvature, we identified two genes whose arabidopsis’s orthologs are involved in stomatal development. They are *cyclin D4*, suggested to control cell division in the initial step of stomata formation in the hypocotyl [56], and *cdc6-B,* an homolog of *cdc6* in arabidopsis, known to be a regulator of stomatal cell division and satellite stomata production [57]. In maize they do not present a putative function yet. For abaxial ACF1 we identified arabidopsis’ *EPF2* (*EPFL1* in maize) and *GAUT10* (not characterised in maize) on chromosome 10 at 143 Mbp. *EPF2*, from the EPFs family, is known to control the stomatal spacing and patterning in the epidermis (Hunt e Gray 2009), while *GAUT10* is important for stomata development and dynamics in combination with *GAUT11* [58], found for aggregation ratio on chromosome 9 at 117 Mbp.

We identified novel QTL associated with vein traits. Among the LVD loci, only the chromosome 2 QTL aligns closely with a region identified by [20] in a field-grown MAGIC maize population. They phenotyped vein density on cross-sections (5^th^ and 9^th^ leaves) using manual dissection, agar embedding, and fluorescent staining. They noted high phenotypic plasticity across different environmental conditions, which might explain why distinct QTL were detected in our dataset. Compared to their labour-intensive histological protocol, our direct surface-imaging approach on ethanol-cleared leaves offers a high-throughput method for vein phenotyping. For LVD, we identified *Arabidopsis’s ERL2* (*ERL3* in maize). EPFs are perceived by receptor complexes in the membrane formed of ERECTA-family receptor like protein kinases [59]. However, this gene is located 6 Mbp from the peak on chromosome 9 and thus its direct involvement in governing LVD remains uncertain. Finally, we identified a single unique QTL for the SD/VD ratio, which co-localises with a QTL for SD on chromosome 6. This genetic overlap supports our hypothesis that stomatal density is the primary driver of the variation in this anatomical ratio and plays a predominant role in modulating plant WUE. Overall, our mapping results support the hypothesis that stomatal and vascular traits are under distinct genetic control at early vegetative stages in maize. Although natural and artificial selection may have shaped the coordination between stomata and veins in domesticated germplasm to balance water supply and transpirational demand, the recombination present in our mapping population allowed us to observe their underlying independent genetic control. The identification of QTL influencing vein density and stomatal density independently represents a major step forward for maize improvement. Our results demonstrate that leaf hydraulic capacity and stomata can potentially be uncoupled and targeted separately. From a breeding perspective, this paves new avenues to manipulate maize stomatal density with no impact on veins, or vice versa. We believe that this evidence can significantly contribute to optimize WUE in a climatic-specific manner.

## Conclusion

Our imaging pipeline combined with QTL mapping approach resulted to be low-cost and high-throughput allowing us to process a high number of samples and images with a discrete image resolution. This pipeline may be a user-friendly support to investigate the same traits on other C4 species, and we could detect other finer stomatal complex features applying a low-cost device with higher resolution. We dissected the stomatal density into a set of stomatal traits and distribution patterns, as well as the vein traits. We identified novel QTL associated with stomatal density, spatial distribution pattern and vein density, suggesting a decoupled contribution to WUE and an independent genetic control of stomatal and vein density. This finding provides a novel genetic framework to independently optimize leaf hydraulic capacity and gas exchange for target environments. Finally, we suggest 10 putative candidate genes in maize involved in stomatal development and patterning, that should be validated with an approach of reverse genetics.

## Availability of Source Code

Stomata and vein detection pipeline

- https://github.com/RPorcedda/Stomatyolo
- Operating system: Platform independent
- Programming language: Python
- Other requirements: see public environment file QTL mapping analysis
- https://github.com/capleo/MAGIC
- Operating system: Platform independent
- Programming language: R
- Other requirements: see public environment file

## Additional Files

Supplementary File S1. Detailed Description of Spatial Distribution Metrics.

Supplementary Table S1. Metadata accession of MAGIC maize lines.

Supplementary Table S2. Description of stomatal and vein traits.

Supplementary Table S3. Broad-sense heritability of the traits.

Supplementary Table S4. QTL peaks across the genome.

Supplementary Table S5. QTL peaks with confidence interval of 10 Mbp across the genome.

Supplementary Table S6. List of candidate genes.

Supplementary Fig. S1. A) Image example from the dataset. B) Stomata annotation on Roboflow. C) Veins detection on Python. D) Detected stomata with height profiles and veins fitted median lines.

Supplementary Fig. S2. Typical spatial distributions of stomata observed inside bundles and relative values of ACF1, Curvature and Flip Rate. ↑ indicates high values, ↓ low values and → medium values.

Supplementary Fig. S3. Fine-Tuning history of pretrained YOLOv8-OBB.

Supplementary Fig. S4. Validation set results: (A) Precision-Recall Curve; (B) F1 score – Confidence; (C) Precision – Confidence; (D) Recall – Confidence. We highlight two confidence thresholds: one for which we obtain the maximum F1 score; one for which we minimize the average difference of stomata count between detected and annotated.

Supplementary Fig. S5. Pearson correlation of abaxial and adaxial leaf side of all stomatal and vein traits.

Supplementary Fig. S6. Pearson correlation matrix on BLUPs values of 273 RILs and 9 founder lines. Correlation coefficient is written on the bottom left and shown by the colour legend, while the significant statistics is shown by the stars on the top right.

Supplementary Fig. S7. Boxplots of BLUPs on six traits of 273 RILs and 9 founder lines. RILs are grey dots while founder lines are coloured.

## Abbreviations

API: application programming interface
BLUP: best linear unbiased predictor
CLAHE: contrast-limited adaptive histogram equalization
DAS: days after sowing
YOLO: You Only Look Once
IoU: intersection over union
mAP: mean average precision
NPK: nitrogen–phosphorus–potassium
PCA: principal component analysis
ProbIoU: probabilistic intersection over union
QTL: quantitative trait locus/loci
SNP: single-nucleotide polymorphism.

## Acknowledgments

We thank Afewerki Yohannes Kiros for developing the experimental design of the greenhouse experiment. We thank Massimo Galbiati, Eleonora Cominelli, and Riccardo Marzali for their support with sample storage. We are also grateful to Giulia Castorina, Gabriellla Consonni and Alberto Caffi for granting access to the greenhouse facilities of the Department of Agricultural and Environmental Sciences at the University of Milan and for their assistance during sampling activities. We further acknowledge Alice Trivellini and Susanna Cialli for sharing laboratory equipment. We also thank Ben Skinner for his input and critical discussions regarding the methodologies employed in this study.

We are also grateful to Svenja Mager, Monique Salardi, Worku Kebede Tekle, Godman Mukamanasasira, Geon Kang, Eshetu Zewdu Tegegn, Mulugeta Tilahun Melaku and Robel Takele Miteku for helping with the annotation process.

## Author Contributions

MS did produce the biological samples and the imaging dataset with the help of MP. RP designed the stomata and veins detection pipeline. MS and RP performed data analyses with inputs from LC, JNF and MDA; MS and RP wrote the paper with input from all the authors. LC, MDA and AV designed the study.

## Data Availability

The imaging data supporting the findings reported in this research are available in the Zenodo Repository https://doi.org/10.5281/zenodo.21770466. The raw DNA sequences from which SNP markers were derived and used in this research are available at the European Nucleotide Archive (ENA) (https://www.ebi.ac.uk/ena/browser/home), under project ID PRJEB67515.

